# Transcriptional plasticity of stromal cells amplifies their differentiation efficiency *in vitro*

**DOI:** 10.1101/2025.04.18.649582

**Authors:** Ali Jasim Mohammad Jamil, Mikkel Ørnfeldt Nørgård, Emilie Grupe, Alexander Rauch

## Abstract

Human bone marrow-derived stromal cells (also termed mesenchymal stem cell -MSCs) are progenitors capable of differentiating into bone forming osteoblasts and fat storing adipocytes. Due to the loss of bone mass being associated with increased marrow fat, trans-differentiation of osteoblasts into adipocytes been hypothesized as a contributor to osteoporotic bone loss and fragility. Reprogramming of transcriptional networks is a prerequisite for cellular differentiation, however, to which extent cell-type specific transcriptional networks modulate cellular plasticity within stromal cells remains unknown. In this study we performed gene expression analysis at bulk and single cell level in stromal cells being repeatedly exposed to osteogenic and adipogenic inducers *in vitro*. Surprisingly, cell type specific gene networks are not suppressive but instead promoting to obtain an opposing phenotype, e.g., enhanced osteoblast differentiation of adipogenic pre-stimulated stromal cells compared to undifferentiated ones. Mechanistically, lineage-selective genes with enhanced response upon interconversion are primed in the stem cell state and obtain modest activity levels during exposure to opposing lineage conditions. Finally, the presence of cells simultaneously showing an osteogenic and adipogenic phenotype highlighting a strong molecular plasticity of transcriptional networks in stromal cells. These observations provide a strong molecular support for the notion of not only progenitor specification but also the plasticity of differentiated cells contributing to the balance of bone mass and marrow fat content.

**Summary:** We provide mechanistic evidence, that cells with established lineage-selective gene programs are more vulnerable to obtain gene signatures and phenotypes of opposing lineages, building ground for trans-differentiation between osteoblasts and adipocytes being a coordinated process rather than random and misdirected.

## Introduction

Bone is a dynamic organ that continuously undergoes remodeling a process initially starting with osteoclasts resorbing old bone from the surface followed by the bone forming actions of osteoblasts. In healthy individuals, there is a balance in bone remodeling meaning that the amount of bone resorption and formation is equal (1, 2). However, during ageing bone mass decreases significantly mainly due to a decrease in bone forming capacity (3) and this decrease in bone mass correlates with an increase in marrow adipocyte volume both during ageing but also in subjects with idiopathic osteoporosis (4–6). This inverse relationship of bone and adipose tissue mass has been at least in part attributed to altered lineage speciation of bone resident stromal progenitor cells that give rise to both osteoblasts and adipocytes (7, 8). In addition, trans-differentiation of mature osteoblasts into adipocytes has been suggested to contribute to the increase of adipocytes on the cost of osteoblasts *in vivo* (9). Here, trans-differentiation describes a process in which cells with an established osteogenic phenotype directly obtain an adipogenic one. The concept of trans-differentiation has not fully been explored in human bone due to lack of lineage tracing techniques. In contrast, genetic mouse models with osteoblast directed overexpression of *Pparg* (10) showed loss of osteoblast function with concordant bone loss while driving expression of a mutated alpha subunit of the stimulatory G protein (Gsa^R201C^) in adiponectin expressing cells leads to loss of adipocytes and increase in bone mass (11). These studies highlight that initial lineage commitment is not definite and can be overcome by strong signals driving maturation into another lineage *in vivo*.

To this day, several *in vitro* studies have demonstrated that stromal cells (MSCs) retain lineage plasticity even after initial commitment. Early work showed that clonally expanded human marrow adipocytes can give rise to osteoblast-like progeny upon subsequent osteogenic stimulation (12), indicating interconversion between adipogenic and osteogenic phenotypes. Likewise, experimental strategies in which stromal cells were first differentiated towards one lineage and subsequently exposed to an alternative induction cocktail demonstrated that osteogenic or adipogenic stimulated cells can adopt chondrogenic, adipogenic or osteogenic phenotypes in response to changed extracellular cues (13–16). However, such findings were considered controversial since other groups reported trans-differentiation to only be the result of progenitor cell contamination or cell fusion during cell culturing (17, 18). Studies that track stromal cells during *in vitro* differentiation clearly showed that cells achieving a mature osteoblast phenotype were capable of trans-differentiating into adipocytes (14). In addition, within mice calvaria individual osteoblasts were shown to adopt an adipogenic fate due to high levels of *Pparg* expression (19). Further, adipogenic committed cells expressing Gsa^R201C^ were capable to form ectopic bone in transplantation assays, supporting the capability of *Adipoq*-expressing stromal cells to undergo bone formation (11).

We recently described that the cellular identity of osteoblast and adipocyte differentiation is determined by lineage-selective transcription factor networks and the confinement of the 3-d-chromatin structure in stromal progenitor cells (20, 21). However, cellular and transcriptional plasticity of human stromal progenitors has not been studied in a way that systematically dissects whether established cell-type specific transcriptional networks oppose or facilitate commitment to other lineages when differentiation conditions are repeatedly switched. Previous trans-differentiation studies primarily focused on single switching events and a limited number of markers (12–14, 16, 22), and therefore did not resolve how many cells resist or succumb to lineage reprogramming, nor how lineage history affects subsequent differentiation capacity and epigenetic responsiveness. In this study we performed histochemical and gene expression analysis at the bulk and single cell level to trace lineage-specific gene networks in immortalized human stromal cells of bone marrow origin (23) that have been exposed to adipogenic or osteogenic inducers in a sequential and alternating manner, respectively. We show a strong plasticity of stromal progenitors, both at the pool and single cell level, where maturing cells were indistinguishable whether they were derived from undifferentiated or trans-differentiated cells. While osteogenic and adipogenic gene programs were opposing each other over time, we propose that single or multiple initiations of cell-type specific gene programs facilitate and enhance the differentiation capacity of stromal cells into opposing lineages. Intriguingly, cell pools that were frequently exposed to short time frames of alternating lineage inducers showed the presence of mature osteoblast and adipocyte phenotypes.

## Results

### Pre-differentiation did not abrogate differentiation ability to opposing lineage

Immortalized human stromal cells from bone marrow (hBMSC-TERT4 cells, (23)) were differentiated *in vitro* using ascorbic acid, b-glycerol phosphate, calcitriol, and dexamethasone for osteogenic induction and insulin, isobutyl methylxanthine, dexamethasone and rosiglitazone for adipogenic induction (fig. 1A). The progression in osteogenic and adipogenic differentiation was followed by the extent of matrix mineralization and lipid droplet formation using staining with Alizarin red and Oil Red’O, respectively. Both matrix mineralization and lipid droplet formation were absent at day 7, established around day 9 to 11, and further increased with prolonged exposure to the differentiation cocktail (fig. 1B). Revisiting RNA-seq data throughout hBMSC-TERT4 cell differentiation (20) we could show that most of the transcriptional reprogramming (represented by PC1) throughout osteoblast or adipocyte differentiation occurred within the first 7 days of differentiation (fig. 1C). However, we could show that only lipid droplet formation occurred in cells that have been withdrawn of lineage inducers after 7 days of differentiation (fig. 1D), while matrix mineralization seemed to be dependent on further osteogenic stimulation. Next, we wanted to test whether hBMSC-TERT4 cells were capable to obtain matrix mineralization upon prior induction along the adipogenic lineage and vice versa. Unexpectedly, we did neither observe a diminished osteogenic nor adipogenic potential when cells were pre-exposed to inducers of the opposite lineage irrespective of exposure time (fig. 1E). Thus, at least at the population level, osteogenic and adipogenic differentiation potential of hBMSC-TERT4 cells was not diminished due to prior commitment towards the opposing lineage.

**Figure 1:**
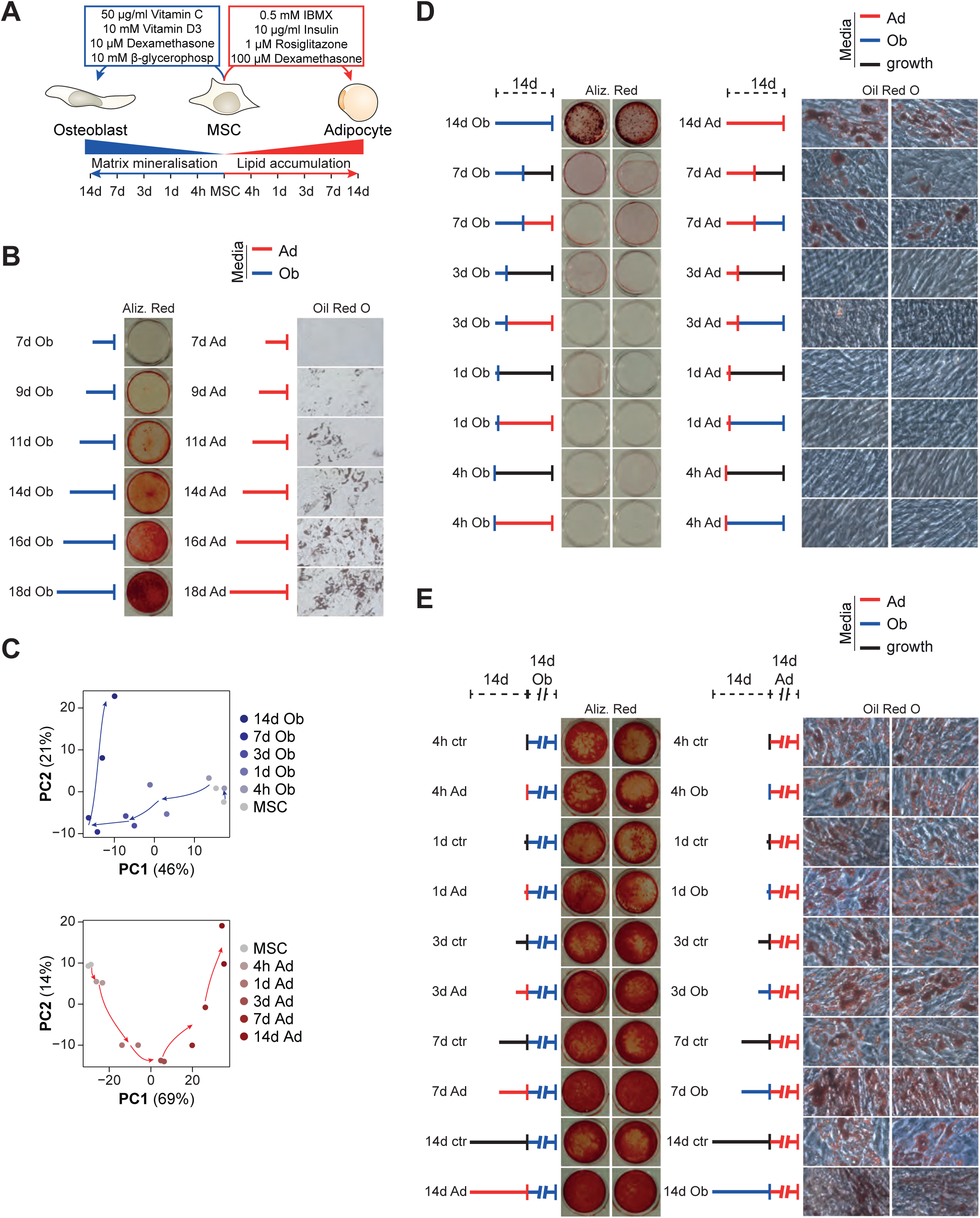
hBMSC-TERT4 cells show phenotypic plasticity. **A)** Model showing differentiation protocol and time points used to harvest RNA for gene expression analyses. **B)** Matrix mineralization (Alizarin Red) and lipid droplet formation (Oil Red’O) in hBMSC-TERT4 cells after the indicated time points of osteoblast (Ob) or adipocyte (Ad) differentiation. **C)** PCA plot from RNA-seq data in osteogenic (upper panel) and adipogenic (lower panel) differentiated hBMSC-TERT4 cells for indicated time points. Arrows are hand drawn for visualization. **D)** Matrix mineralization (left panel) and lipid droplet formation (right panel) of hBMSC-TERT4 cells after 14 days of differentiation initiation. Cells were initiated for the indicated time points and switched to growth media (black), osteogenic media (Ob -blue), or adipogenic media (Ad -red) for the remaining time. **E)** Matrix mineralization (left panel) and lipid droplet formation (right panel) in hBMSC-TERT4 cells exposed to 14 days of osteogenic and adipogenic cocktail after pre-stimulation with growth media (black), osteogenic media (Ob -blue), or adipogenic media (Ad -red) for the indicated time points.

### hBMSC-TERT4 cells showed great plasticity at the transcriptional level upon switching lineage inducers

We performed bulk RNA-sequencing on hBMSC-TERT4 cells committing towards the osteogenic or adipogenic lineage for 7 days followed by being exposed to the opposing inducers for 4 days (fig. 2A). Focusing on putative osteogenic or adipogenic marker as well as stem-cell associated genes (fig. 2B), dimensionally reduction of global gene expression by principle component analysis (fig. 2C), or previously defined (20) osteoblast- and adipocyte-selective gene signatures (fig. 2D), we could demonstrate the obliteration of the initial and establishment of the new lineage-selective gene programs upon switching differentiation cocktails in hBMSC-TERT4 cells.

**Figure 2:**
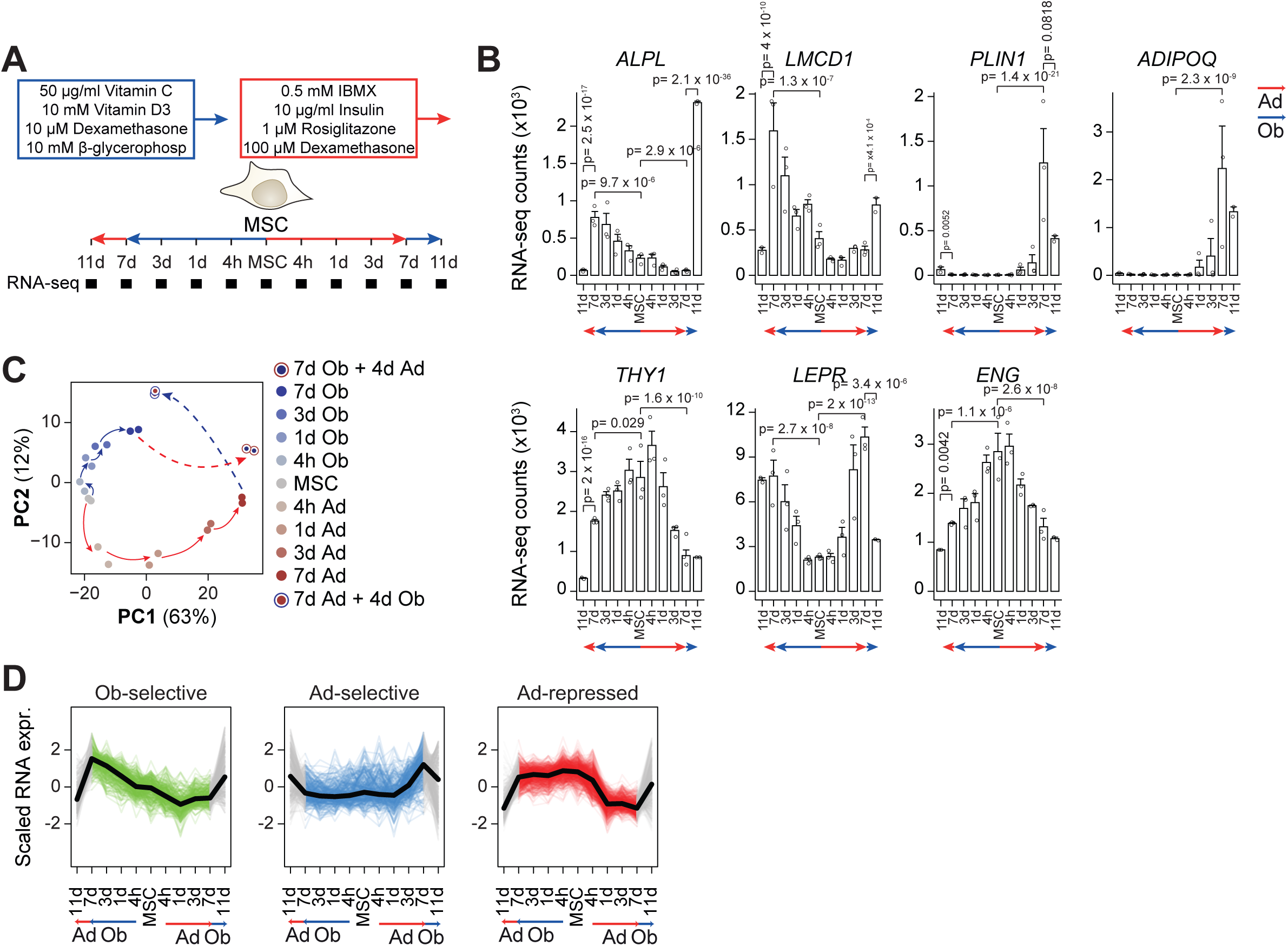
hBMSC-TERT4 cells show transcriptional plasticity. **A)** Model showing differentiation protocol and time points used to harvest RNA for gene expression analyses. **B)** Bar plot of RNA-seq derived gene counts of osteoblast-specific *ALPL* and *LMCD1*, adipocyte-specific *PLIN1* and *ADIPOQ*, and stem cell-associated *THY1*, *LEPR*, and *ENG* expression. DESeq2 derived adjusted p-values are indicated for p < 0.1 with focus on 7 days and switching conditions. **C)** PCA plot from RNA-seq data of hBMSC-TERT4 cells for indicated time points. Arrows are hand drawn for visualization. **D)** Line plots showing scaled gene expression levels of previously identified gene signatures that are osteoblast-selective induced (n = 291), adipocyte-selective induced (n = 358), or adipocyte-selective repressed (n = 766). Black line indicates mean. Colored lines show normal differentiation, and grey lines show media switching.

### scRNA-seq analysis indicated similar lineage commitment from undifferentiated or pre-differentiated cells

Since hBMSC-TERT4 cells showed great plasticity for osteogenic and adipogenic gene expression at the population level we generated scRNA-seq libraries from undifferentiated, 7 days committed, and 7 days + 4days trans-differentiated cells (fig. 3A). We questioned whether cells exhibit expression of both osteoblast and adipocyte-specific genes and whether transcriptional reprogramming differs in cells upon direct lineage commitment versus trans-differentiation. UMAP dimensionality reduction distinguished cells of the five libraries according to treatment protocol (fig. 3B), and scRNA-seq libraries showed similar dynamics for osteoblast-selective, adipocyte-selective, and stem cell-associated markers compared to bulk RNA-seq (fig. 3C). In line with the results of the PCA from our bulk RNA-seq experiments (fig. 2C), we observed a close relationship of the scRNA-seq libraries according to the final differentiation cocktail. Interestingly, using the expression of osteoblast and adipocyte selective gene signatures ((20), fig 3D) as well as pseudotime analysis (fig. 3E and arrows in 3B) showed convergence of highly differentiated cells. This indicates that cells which committed towards osteoblasts or adipocytes from either an un- or pre-differentiated state are almost indistinguishable at the single cell level. In addition, scRNA-seq enabled us to identify cells that did not respond to the addition of the opposing inducers, i.e., cells expressing lineage-selective genes according to the initial differentiation cocktail (fig. 3F). Here, we detected cells that maintained a strong adipogenic signature despite switching to osteogenic inducers (7D Ad + 4d Ob, fig. 3F second panel), which is in line with the observation of cells with lipid-filled droplets in cultures that received 7 days adipogenic followed by 7 days osteogenic stimulation (fig. 1D).

**Figure 3:**
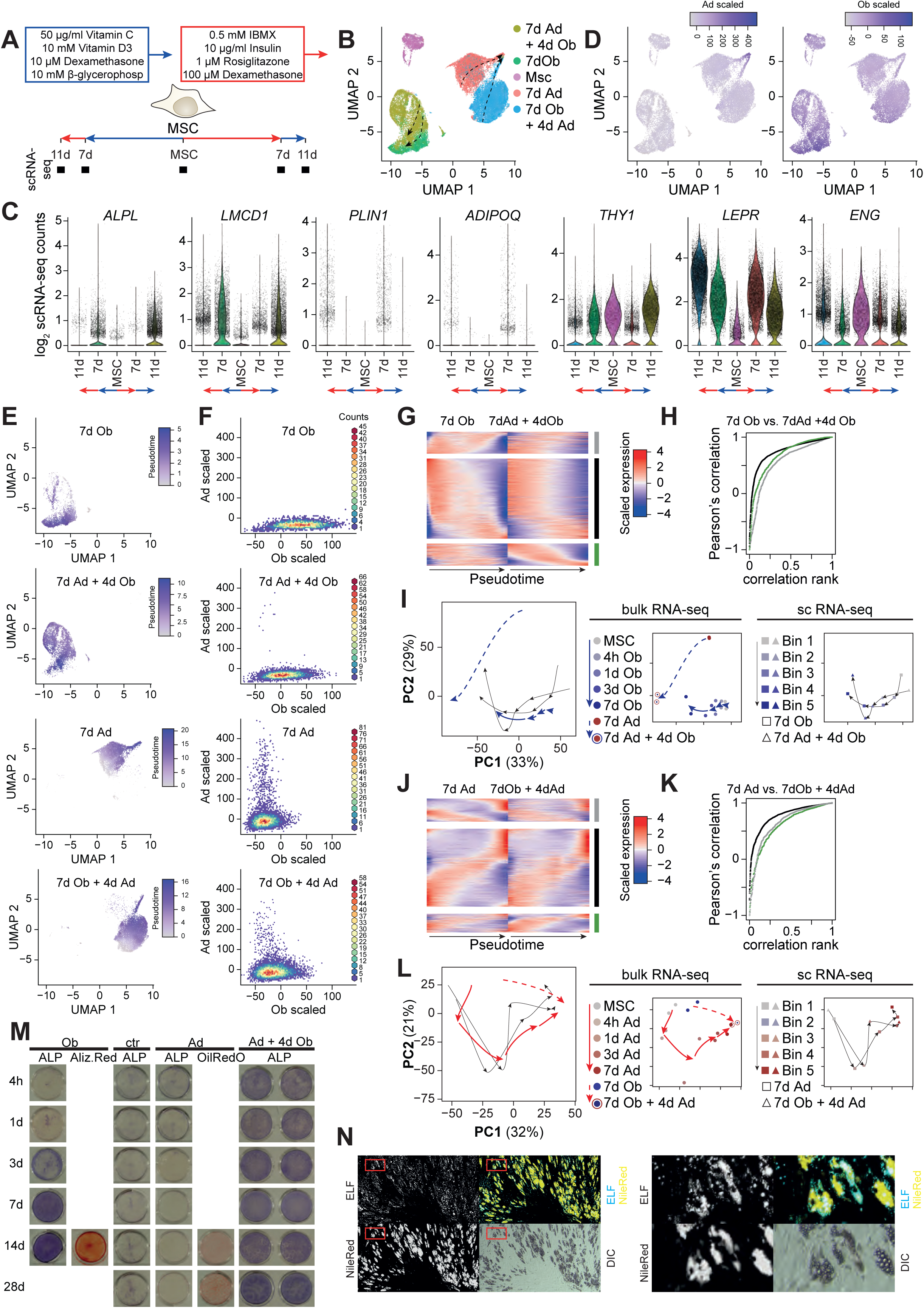
hBMSC-TERT4 cells show lineage plasticity at single cell level. **A)** Model showing differentiation protocol and time points used to harvest RNA for single cell gene expression analyses. **B)** UMAP-plot showing scRNA-seq libraries of hBMSC-TERT4 cells. Arrows indicate pseudotime trajectories for individual samples. **C)** Violin plots denoting expression of osteoblast markers *ALPL* and *LMCD1*, adipocyte markers *PLIN1* and *ADIPOQ*, and stem cell-associated markers *THY1*, *LEPR*, and *ENG* across scRNA-seq samples of hBMSC-TERT4 cells. **D)** Feature plot quantifying summed expression levels of previously defined gene signatures that are adipocyte-specific induced (upper panel) or osteoblast-specific induced (lower panel). Sum was calculated using scaled expression levels. E) Feature plot showing pseudotime values of individual RNA-seq samples. These data are the basis for the arrows in fig. 3B. **F)** Density plot contrasting summed expression levels of previously defined gene signatures that are adipocyte-specific (Ad-scaled) and osteoblast-specific (Ob-scaled) induced for the individual scRNA-seq samples. **G)** Heat map showing genes differentially expressed (p < 0.001) and ordered by pseudotime (100 bins) analysis in hBMSC-TERT4 cells receiving finally osteogenic media. Grey, significant in 7d Ob, green significant in 7d Ad + 4d Ob, black significant in both samples. **H)** Ranked Pearson’s correlation of pseudotime expression in 7d Ob and 7d Ad + 4d Ob of gene groups in fig. 4G. **I)** PCA plot combining trajectories of bulk RNA-seq (small panel left) and scRNA-seq (small panel right) for genes with differential expression (p < 0.001) during pseudotime analysis in hBMSC-TERT4 cells receiving finally osteogenic media (fig. 4G). scRNA-seq data were averaged across 5 bins and centered based on the PCA environment of the bulk RNA-seq data. **J)** Heat map showing genes differentially expressed (p < 0.001) and ordered by pseudotime (100 bins) analysis in hBMSC-TERT4 cells receiving finally adipogenic media. Grey, significant in 7d Ad, green significant in 7d Ob + 4d Ad, black significant in both samples. **K)** Ranked Pearson’s correlation of pseudotime expression in 7d Ad and 7d Ob + 4d Ad of gene groups in fig. 4J. **L)** PCA plot combining trajectories of bulk RNA-seq (small panel left) and scRNA-seq (small panel right) for genes with differential expression (p < 0.001) during pseudotime analysis in hBMSC-TERT4 cells receiving finally adipogenic media (fig. 4K). scRNA-seq data were averaged across 5 bins and centered based on the PCA environment of the bulk RNA-seq data. **M)** hBMSC-TERT4 cells were either maintained in culture media (ctr), osteogenic media (Ob), or adipogenic media (Ad) for the indicated time points (rows) and if indicated treated with additional 4 days of osteogenic media (columns). ALP activity, matrix mineralization (Alizarin Red) or lipid accumulation (Oil Red’O) was assessed (columns). **N)** hBMSC-TERT4 cells were exposed to adipogenic inducers for 28 days followed by 4 days of osteogenic stimulation (corresponds to lower right pictures in fig. 4M). Lipid droplet formation and ALP activity was determined by NileRed (yellow) or ELF fluorescence (Blue) staining. 10 x magnification, DIC -Differential Interference Contrast.

To get a deeper understanding of the transcriptional remodeling when comparing cells that undergo direct differentiation and cells that show strong plasticity when exposed to switching differentiation cocktails, we compared gene expression dynamics along pseudotime in our scRNA-seq data. Focusing on cells that finally received the osteogenic differentiation cocktail (7dOb and 7dAd + 4dOb), we could identify three groups of genes with differential expression (expressed in 10% of cells and p-val < 0.001) along pseudotime, 1) only in direct differentiated cells (7dOb, fig. 3G grey group), 2) only in cells with switched media exposure (7dAd + 4dOb, fig. 3G green group), and 3) in both (fig. 3G black group). However, for all three groups we could see a strong correlation of gene expression in both scRNA-seq libraries (fig. 3H), indicating differences in p-value thresholds rather than actual dynamics. For further confirmation we could show that except for 14 days of osteoblast differentiation bulk expression data recapitulated the pseudotime-based expression dynamics (fig. 3I). This demonstrates that at least for the time points investigated, osteogenic commitment from undifferentiated and adipogenic stimulated cells followed almost identical transcriptional reprograming. On the other side, we made similar observations for adipogenic commitment from undifferentiated and osteogenic differentiated cells (fig. 3J – L).

Given the fact that almost all cells of a differentiating pool showed strong transcriptional plasticity and that transcriptional remodeling during direct and trans-differentiation followed similar principles we hypothesized that cells which initiate a lineage-specific gene program are also likely to express genes related to the other lineage when induction cocktails are switched. To test this, we followed ALP activity along the adipogenic lineage for up to 28 days followed by a stimulation with osteogenic media for 4 days (fig. 3M). Induction of ALP activity was not diminished as more and more cells accumulated lipid droplets during the course of pre-differentiation towards the adipogenic lineage. Next, we employed fluorescent staining on ALP activity and lipid droplet formation in hBMSC-TERT4 cells that have been exposed to 28 days of adipogenic cocktail followed by 4 days of osteogenic one. Intriguingly, we observed many adipocytes based on Nile Red staining (fig. 3N) and exactly those areas (fig. 3N, upper panel) and cells (fig. 3N, lower panel) occupied by lipid filled droplets showed strong induction of ALP activity. This affirmed our hypothesis that cells capable to form lipid droplets are also able to activate ALP activity upon osteogenic stimulation.

### Pre-commitment potentiated the differentiation potential of stromal cells

Having shown that hBMSC-TERT4 cells have an enormous plasticity in terms of differentiation quality we next questioned whether pre-exposition to adipogenic inducers enhances osteogenic potential and vice versa. Adipogenic and, although to a lesser extent, osteogenic pre-stimulation enhanced matrix mineralization (fig. 4A) and lipid droplet formation (fig. 4B) of cultures that have been switched to osteogenic and adipogenic inducers, respectively. These observations were supported by gene expression analysis of the osteoblast and adipocyte-selective genes *ALPL* and *ADIPOQ* (fig. 4C). Next, we wanted to test whether this enhanced osteogenic or adipogenic potential due to opposing pre-differentiation could rescue bio-banked primary cells that have been previously identified with low osteogenic or adipogenic potential (24). Like the observations in hBMSC-TERT4 cells we could show a strong enhancement of the osteogenic potential if cells have been pre-exposed to adipogenic inducers (fig. 4D & E), with enabling mineralization in cultures that did otherwise not show any signs of late osteoblast commitment during directed differentiation. On the other side, enhancement of adipogenic differentiation due to prior osteogenic differentiation was mild (fig. 4F) in line with observations from hBMSC-TERT4 cells. Finally, we tested whether pre-stimulation with adipogenic inducers *in vitro* will change the bone forming potential of hBM-MSC-TERT4 cells *in vivo* using ectopic transplantation assays into immune deficient mice. Testing hBM-MSC-TERT4 sublines with low and high bone forming potential (25), we observed an increase in bone in implants of bone-forming competent cells pretreated with adipogenic inducers (fig. 4G & H). In contrast, cells not capable of bone formation did neither form bone if exposed to adipogenic inducers nor control media (fig. 4H). These results collectively show that adipogenic and osteogenic lineage specification *in vitro* potentiates the differentiation potential of stromal progenitors from bone marrow origin towards osteoblasts and to a lesser extent towards adipocytes, respectively.

**Figure 4:**
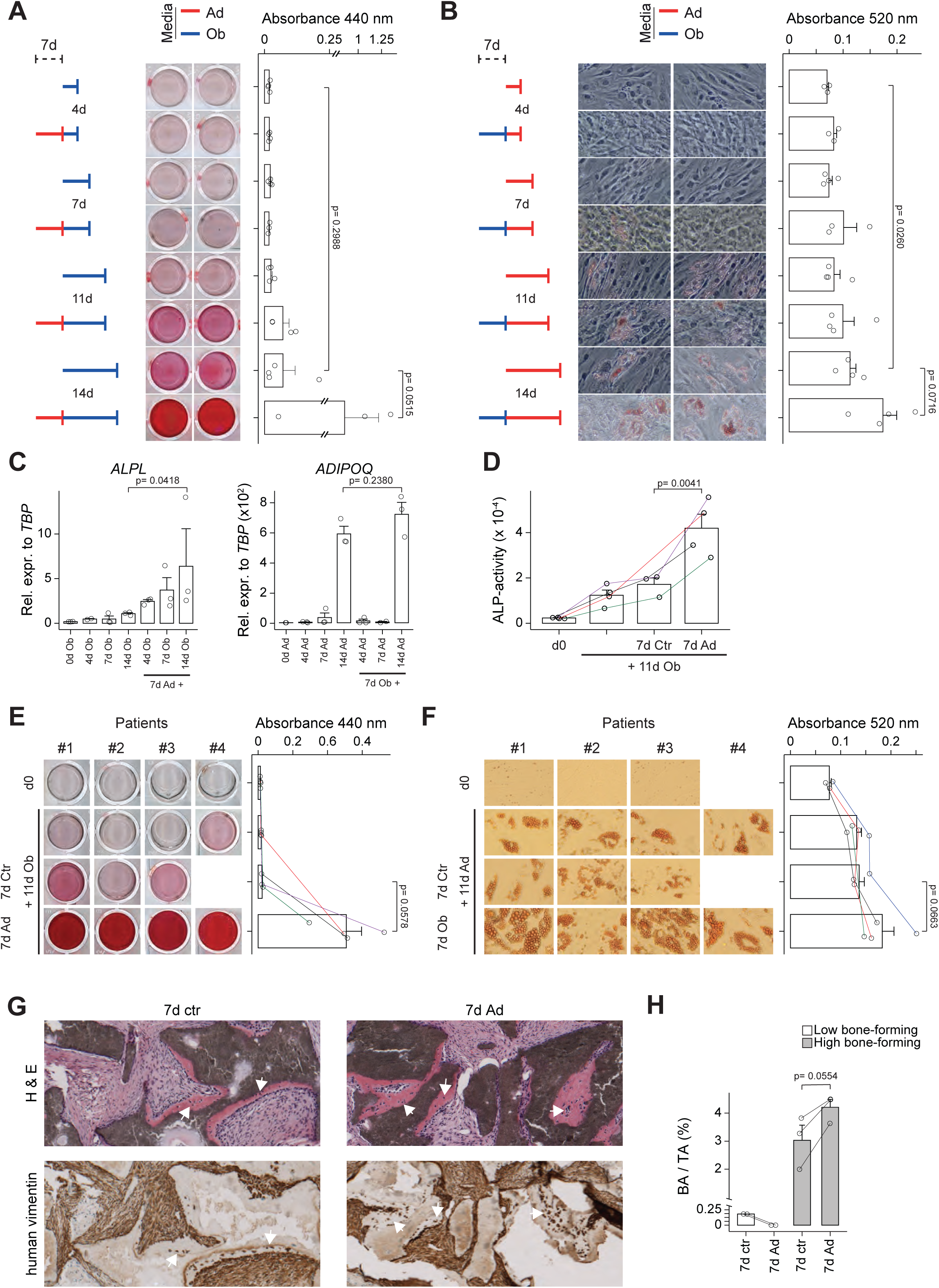
Lineage switching amplifies differentiation potential of stromal cells. **A)** hBMSC-TERT4 cells were directly differentiated for the indicated time points to the osteogenic lineage (Ob – blue) or pre-treated with adipogenic inducers (Ad – red) for 7 days. Matrix mineralization (Alizarin Red) was assessed and quantified. Unpaired t-test, n= 3. **B)** hBMSC-TERT4 cells were directly differentiated for the indicated time points to the adipogenic lineage or pre-treated with osteogenic inducers for 7 days. Lipid droplet formation (Oil Red’O) was assessed and quantified. Unpaired t-test, n= 4. **C)** Relative mRNA expression levels of *ALPL* (left panel, cells from fig. 4A) and *ADIPOQ* (right panel, cells from fig. 4B) in hBMSC-TERT4 cells. Unpaired t-test, n= 3. Expression was normalized to *TBP*. **D)** Biobanked primary stromal cells were directly differentiated for 11 days with osteogenic media or pre-treated for 7 days with culture media or adipogenic inducers. Matrix mineralization (Alizarin Red) was assessed and quantified. Paired t-test, n= 4. Empty panels represent detached cells. **E)** ALP activity was determined in cells from fig. 4D. Paired t-test, n= 4. **F)** Biobanked primary stromal cells were directly differentiated for 11 days with adipogenic media or pre-treated for 7 days with culture media or osteogenic inducers. Lipid droplet formation (Oil Red’O) was assessed and quantified. Paired t-test, n= 4. Empty panels represent detached cells. G) H & E staining (upper panel) and immune staining against human-specific vimentin (lower panel) of close-by sections from ectopic bone formation assays in immune deficient mice. Bone forming competent hBM-MSC-TERT4 cells were pretreated with adipogenic inducers or growth media *in vitro* for 7 days prior to implantation. White arrowheads indicate areas of bone formation with deposition of vimentin positive osteocytes. H) Bar plot quantifying bone as percentage of bone area over tissue area from ectopic transplantation assays of hBM-MSC-TERT4 cells with low or high bone forming capacity into immune deficient mice. hBM-MSC-TERT4 cells were pretreated with adipogenic inducers or growth media *in vitro* for 7 days prior to implantation. Lines connect implants from the same mouse. Paired t-test, n = 2 for low bone forming cells, n = 3 for high bone forming cells.

### hBMSC-TERT4 cells showed repeated transcriptional plasticity

We were intrigued by the transcriptional plasticity of stromal progenitor cells upon switching lineage inducers once and questioned if this plasticity persists after repeated switching. Using bulk RNA-seq in hBMSC-TERT4 cells that switched lineage inducers twice, at day 3 and 7 of differentiation (fig. 5A), we showed that stromal progenitors maintain lineage plasticity at the transcriptional level globally (fig. 5B), for lineage selective gene signatures (fig. 5C), and for selected lineage and stem cell gene markers (fig. 5D). Thus, even cells with increased osteogenic potential due to pre-induction with adipogenic inducers maintained the capacity to re-activate adipogenic gene expression, and vice versa. Using activity (fig. 5E) and mRNA expression (fig. 5F) of alkaline phosphatase, we could demonstrate the ability of cells to enhance and repress an osteogenic phenotype when cells were exposed to osteogenic or adipogenic inducers. Interestingly, repeated changing of lineage inducers built-up both ALP activity and *ALPL* mRNA expression, indicating that at least for osteogenic differentiation alternating lineage commitment increased the responsiveness of cells to osteogenic inducers. Having shown that exposure to adipogenic inducers diminished the osteogenic phenotype but also increased osteogenic responsiveness, at least for alkaline phosphatase, we questioned if a continuous alternating exposure to osteogenic and adipogenic inducers would finally lead to calcification and lipid droplet formation. hBMSC-TERT4 cells exposed to 4 day turns of osteogenic or adipogenic inducers for a total of 32 days (fig. 6A) showed both the presence of lipid droplet filled cells and strong matrix mineralization (fig. 6B). Quantifying mRNA expression of *ALPL* and *ADIPOQ* after 6 to 8 rounds of media switching showed that cells maintained oscillating expression patterns of induction and repression, and that inducibility of *ALPL* and *ADIPOQ* still increased over time (fig. 6C). Using fluorescence microscopy, we could show that ALP activity was especially strong in cells with lipid accumulation (fig. 6D-F) highlighting that these cells despite their adipogenic phenotype had a very strong osteogenic inducibility. Single nuclei sequencing of hBMSC-TERT4 cells subjected to 8 rounds of media switching demonstrated four distinct cell clusters (fig. 6G & H), 1) proliferating cells (cluster 0) strongly expressing genes such as *MDM2*, *ZMAT3* and *SLC7A11*; 2) cells expressing high levels of pre-adipocyte markers (cluster 1) such as *PDGFRA*, *PDGFRB*, and *FGF10*; 3) cells expressing mature adipocyte genes (cluster 2) such as *ADIPOQ*, *PLIN1* and *HSD11B1*; and 4) cells expressing mature osteoblast genes (cluster 3) such as *POSTN*, *RUNX2* and *COL1A1*. The latter two clusters were low in numbers, but interestingly, cells of cluster 3, that express mature osteoblast genes were also high in *PPARG* expression (fig. 6H) and our previously defined adipocyte gene signature (fig. 6I). Quantifying those osteoblast and adipocyte gene signatures at single nuclei level showed, that there were indeed cells expressing both osteoblast and adipocyte associated genes at the same time (fig. 6J). These data indicate that repeated switching of lineage-inducers led to individual cells expressing features from both lineages in line with the high ALP activity levels in lipid-droplet positive cells.

**Figure 5:**
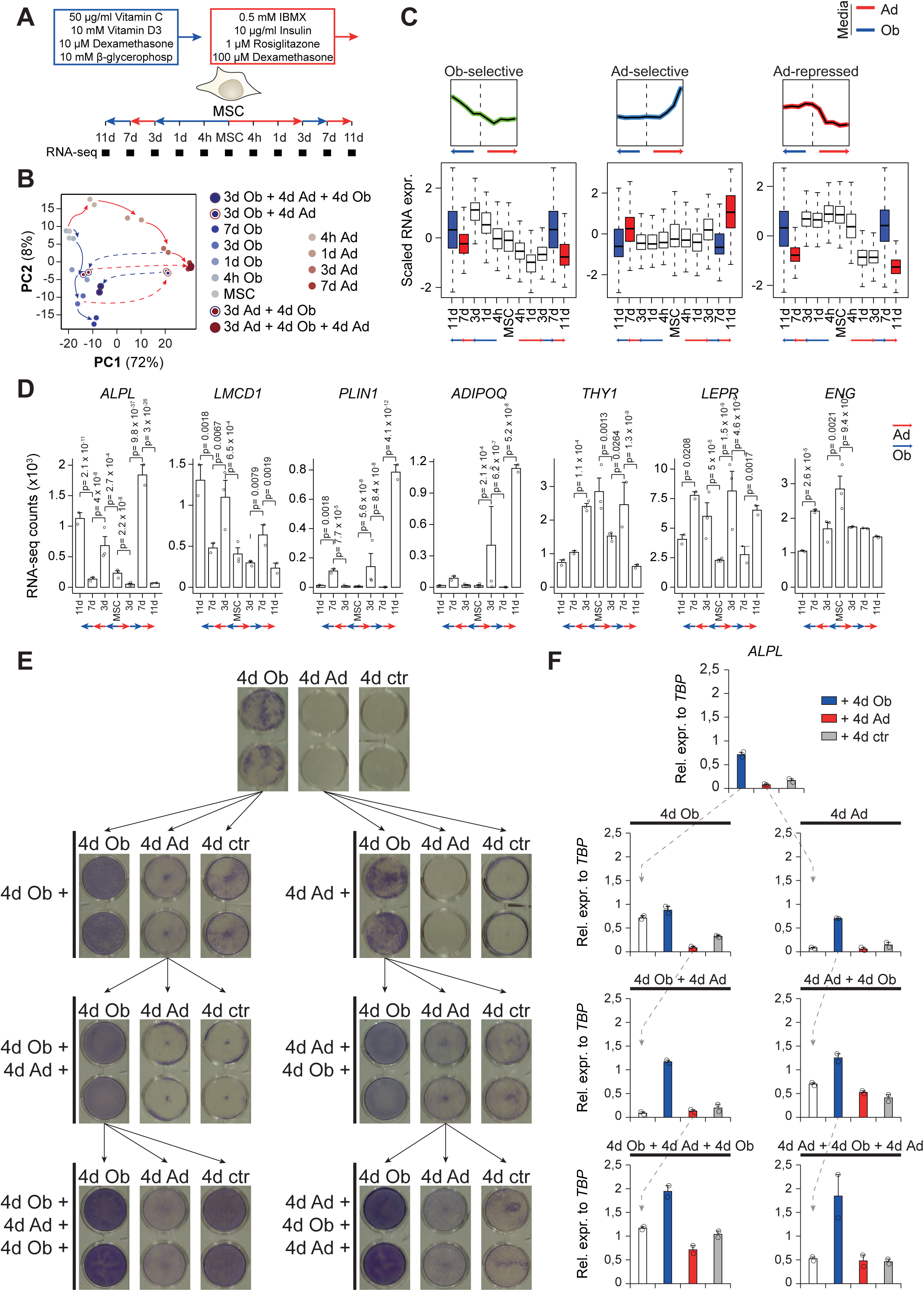
hBMSC-TERT4 cells maintain plasticity after repeated lineage switching. **A)** Model showing differentiation protocol and time points used to harvest RNA for gene expression analyses. **B)** PCA plot from RNA-seq data of hBMSC-TERT4 cells for indicated time points. Arrows are hand drawn for visualization. **C)** Box plot showing scaled gene expression levels of previously identified gene signatures that are osteoblast-selective induced (top, n = 291), adipocyte-selective induced (middle, n = 358), or adipocyte-selective repressed (bottom, n = 766). White boxes indicate direct differentiation; blue boxes change to osteogenic and red box change to adipogenic media. Small graphs to the side show mean expression levels of gene signatures as previously defined (20). **D)** Bar plot of RNA-seq derived gene counts of osteoblast-specific *ALPL* and *LMCD1*, adipocyte-specific *PLIN1* and *ADIPOQ*, and stem cell-associated *THY1*, *LEPR*, and *ENG* expression in hBMSC-TERT4 cells. DESeq2 derived adjusted p-values are indicated for p < 0.1. **E)** ALP staining on hBMSC-TERT4 cells were exposed to culture media (ctr), osteogenic media (Ob), or adipogenic media (Ad) for 4 days and media was afterwards switched along the arrows of the tree up to a total of 4 switches and 16 days stimulation in total. **F)** Relative mRNA expression levels of *ALPL* from cells in Fig 5E. n = 2. White bars correspond to blue or red bars in the panel above. *ALPL* was normalized to *TBP*.

**Figure 6:**
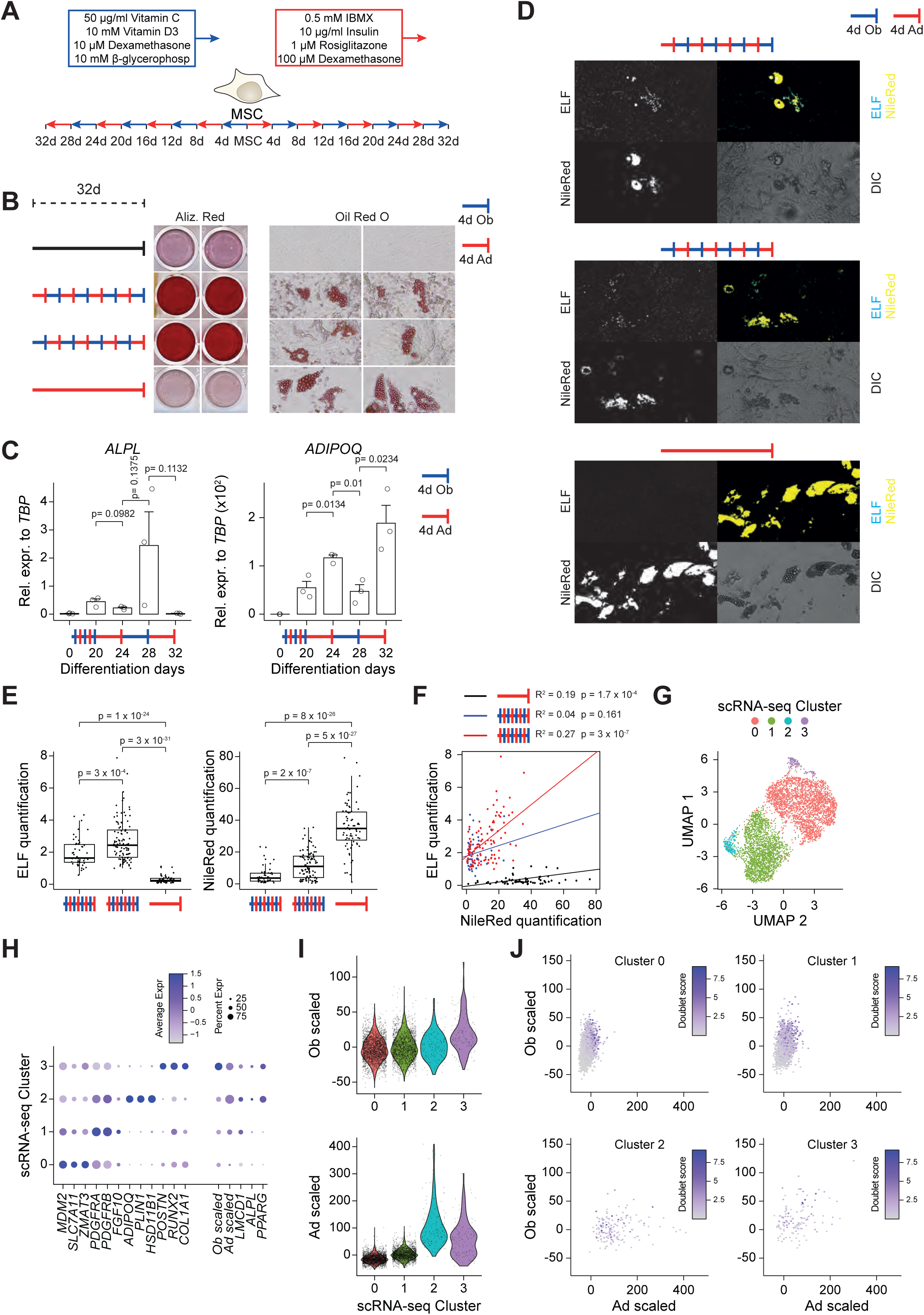
Repeated time-limited lineage switching induces lipid accumulation in calcified cultures. **A)** Model showing differentiation protocol with switching lineage inducers every 4 days for a total of 32 days. **B)** Matrix mineralization (Alizarin Red) and lipid droplet formation (Oil Red’O) in cells maintained in culture media (black) or exposed to indicated differentiation cocktails. **C)** Relative mRNA expression levels of *ALPL* (left panel, cells from fig. 6B third row) and *ADIPOQ* (right panel, cells from fig. 6B second row) in hBMSC-TERT4 cells. Unpaired t-test, n= 3. Expression was normalized to *TBP*. **D)** Lipid droplet formation and ALP activity was determined by NileRed (yellow) or ELF fluorescence (Blue) staining. Upper panel corresponds to cells in fig. 6B second row, middle panel to cells in fig. 6B third row, and lower panel to cells in fig.6B fourth row. 10 x magnification, DIC -Differential Interference Contrast. **E)** Box plot quantifying ELF and Nile Red fluorescence intensity using Nile Red objects of cells shown in fig. 6D. Number of quantified cells when subjected to media switching starting with 1) osteogenic media (n=50), 2 adipogenic media (n=99) or 3) induced along the adipogenic lineage only (n=72). Unpaired t-test. **F)** Scatter plot contrasting quantification of ELF and Nile Red fluorescence intensity at Nile Red spots for cells in fig. 6D. Lines represent linear regression and its statistics are indicated. **G)** UMAP plot of single nuclei RNA-seq of hBMSC-TERT4 cells undergoing differentiation for 32 days, starting with adipogenic stimulation and switching differentiation media every fourth day (corresponds to cells in fig. 6B second row). **H)** Dot plot showing cluster specific gene expression of genes with differential expression across clusters (first 12 genes), previously defined gene signatures that are either osteoblast-selective (Ob scaled) or adipocyte-selective (Ad scaled) induced, *LMCD1*, *ALPL*, and *PPARG* in snRNA-seq of cells in fig. 6G **I)** Violin plots quantifying Ob-scaled (upper panel) or Ad-scaled (lower panel) in clusters of snRNA-seq of cells in fig. 6G. **J)** Scatter plot contrasting Ob-scaled and Ad-scaled in clusters of snRNA-seq of cells in fig. 6G. Color code shows predicted doublet score of the individual nuclei.

### Lineage switching is linked to distinct epigenetic signatures at osteogenic and adipogenic genes

Previous work has shown that enhancer activation is highly dynamic in response to switching lineage inducers in mouse stromal cells *in vitro* (26). Building on this, we asked whether epigenetic signatures differ between genes that are strongly induced and those that fail to be induced upon switching lineage inducers. To address this, we integrated published enhancer activity marks (20) with bulk RNA-seq data from day 0 and 3 of normal differentiation as well as bulk RNA-seq data from cells exposed to opposing lineage inducers for 4 days following 3 days of initial specification (fig. 7A). Enhancers were associated with genes based on genomic proximity. We started with genes activated during adipogenic differentiation and first classified whether those genes were also activated during osteogenic stimulation (Common) or not (Spec). Second, we further divided each group according to whether their induction upon switching osteogenic pre-stimulated cells to adipogenic conditions was higher, similar, or lower compared to normal adipogenic differentiation (fig. 7B). Since loci of commonly activated genes are expected to undergo enhancer activation in both contexts, we focused to distinguish loci of adipocyte-specific genes that showed higher induction levels such as *ABCA9* from those with less such as *PNMA2* (fig. 7C & D). Despite being clearly distinct from commonly induced genes, loci from genes that were more strongly induced upon switching from osteogenic to adipogenic conditions showed modestly increased chromatin accessibility and MED1 recruitment during osteogenic pre-stimulation compared to loci of less induced genes (fig. 7E). These loci did not differ in basal levels of enhancer activity in the undifferentiated state (fig. 7F), suggesting that enhanced adipogenic gene expression is linked to licensing of enhancers during osteogenic pre-stimulation, thereby potentiating their subsequent activation under adipogenic conditions.

**Figure 7:**
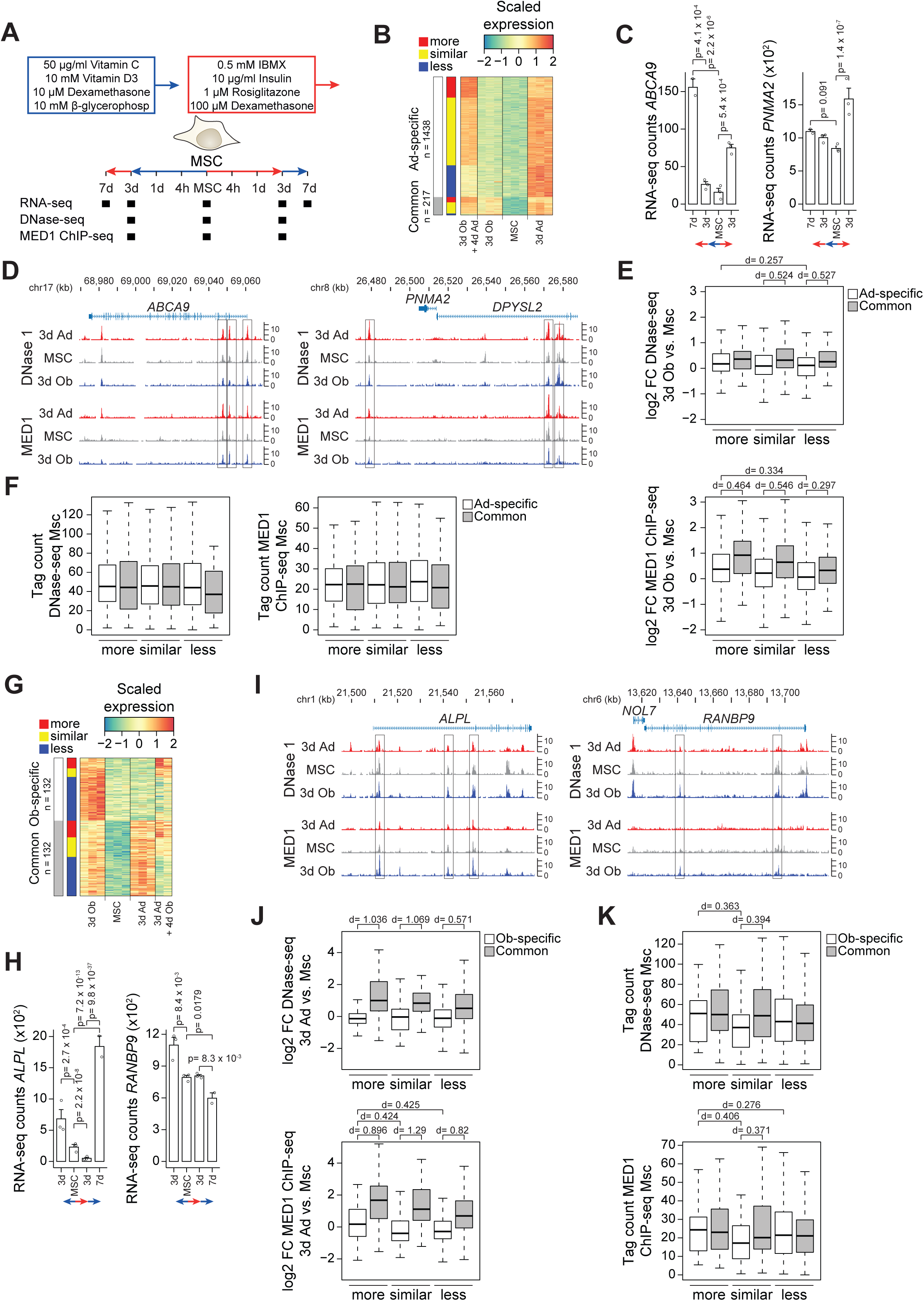
Distinct epigenetic signatures characterize interconversion-responsive osteogenic and adipogenic genes. **A)** Model showing differentiation protocol and time points used to harvest RNA for gene expression analyses as well as DNase-seq and MED1 ChIP-seq for enhancer dynamics. **B)** Heatmap focusing on genes induced during adipocyte differentiation (p-adj < 0.01 & logFC > 0 for 3dAd vs MSC). Genes were grouped into commonly upregulated ones (p-adj < 0.05 & logFC > 0 for 3dOb vs. MSC) and those specifically upregulated during adipocyte differentiation. Subgrouping into more, similar, or less induced was based on k-means clustering of expression data from MSC, 3dAd, and 3dOb + 4dAd. **C)** Bar plot of RNA-seq derived gene counts of adipocyte-specific induced genes with higher, *ABCA9*, and lower induction levels, *PNMA2*, after interconversion of hBMSC-TERT4 cells. DESeq2 derived adjusted p-values are indicated for p < 0.05. **D)** Genome browser screenshots showing tag density of DNase-seq and MED1 ChIP-seq in hBM-MSC-TERT4 cells prior to and after 3 days of osteogenic or adipogenic differentiation. Left panel, loci of *ABCA9* representing an interconversion more induced adipocyte-specific gene. Right panel, loci of *PNMA2* representing an interconversion less induced adipocyte-specific gene. **E)** Box plot quantifying changes in DNase-seq (upper panel) and MED1 ChIP-seq (lower panel) during 3 days of osteogenic differentiation in hBM-MSC-TERT4 cells at MED1 bound enhancers nearby genes grouped in fig. 7B. Effect sizes were calculated using Cohen’s d. **F)** Box plot quantifying DNase-seq (left panel) and MED1 ChIP-seq (right panel) tag density in undifferentiated hBM-MSC-TERT4 cells at MED1 bound enhancers nearby genes grouped in fig. 7B. Effect sizes were calculated using Cohen’s d. **G)** Heatmap focusing on genes induced during osteoblast differentiation (p-adj < 0.01 & logFC > 0 for 3dOb vs MSC). Genes were grouped into commonly upregulated ones (p-adj < 0.05 & logFC > 0 for 3dAd vs. MSC) and those specifically upregulated during osteoblast differentiation. Subgrouping into more, similar, or less induced was based on k-means clustering of expression data from MSC, 3dOb, and 3dAd + 4dOb. **H)** Bar plot of RNA-seq derived gene counts of osteoblast-specific induced genes with higher, *ALPL*, and lower induction levels, *RANBP9*, after interconversion of hBMSC-TERT4 cells. DESeq2 derived adjusted p-values are indicated for p < 0.05. **I)** Genome browser screenshots showing tag density of DNase-seq and MED1 ChIP-seq in hBM-MSC-TERT4 cells prior to and after 3 days of osteogenic or adipogenic differentiation. Left panel, loci of *ALPL* representing an interconversion more induced osteoblast-specific gene. Right panel, loci of *RANBP9* representing an interconversion less induced osteoblast-specific gene. **J)** Box plot quantifying changes in DNase-seq (upper panel) and MED1 ChIP-seq (lower panel) during 3 days of adipogenic differentiation in hBM-MSC-TERT4 cells at MED1 bound enhancers nearby genes grouped in fig. 7G. Effect sizes were calculated using Cohen’s d. **K)** Box plot quantifying DNase-seq (left panel) and MED1 ChIP-seq (right panel) tag density in undifferentiated hBM-MSC-TERT4 cells at MED1 bound enhancers nearby genes grouped in fig. 7G. Effect sizes were calculated using Cohen’s d.

We then reciprocally grouped genes activated during osteogenic differentiation (fig. 7G) and focused on loci of osteoblast-specific genes that exhibited higher or lower induction levels upon switching adipogenic pre-stimulated cells to osteogenic conditions, such as *ALPL* and *RANBP9* (fig. 7H & I). Similar to the osteogenic-to-adipogenic switch, we observed elevated levels of chromatin accessibility and MED1 recruitment upon adipogenic pre-stimulation comparing genes with higher and lower induction levels (fig. 7J). This difference was due to a slight increase in enhancer activity near genes with higher induction levels and a decrease near genes with similar or lower induction levels. In addition, these loci differed in basal enhancer activity in the undifferentiated state, with high accessibility and MED1 recruitment to genes with higher induction levels (fig. 7K). Thus, enhanced osteogenic gene expression following adipogenic pre-stimulation is linked to stem cell-primed enhancers that do not lose activity during adipogenic stimulation. Collectively, our results demonstrate that enhanced lineage-specific gene induction differs between adipogenic-to-osteogenic and osteogenic-to-adipogenic transitions, with distinct genomic loci and enhancer activity states, suggesting differential engagement of regulatory machinery depending on lineage history.

## Discussion

### Stromal cells showed both cellular and transcriptional plasticity upon changing lineage inducers

The aim of this study was to investigate the cellular and transcriptional plasticity of human stromal progenitors towards the osteogenic and adipogenic lineage *in vitro*. We observed high cellular and transcriptional plasticity within hBMSC-TERT4 cells since *ALPL* expression, matrix mineralization, *ADIPOQ* expression and lipid droplet formation were similar or higher in cells exposed to opposing lineage inducers compared with directly differentiated cells. A previous study showed that mature osteoblasts, selected through an osteocalcin promoter driven GFP reporter, were able to obtain both an adipogenic or chondrogenic phenotype (14), indicating maintenance of multipotency in differentiated cells *in vitro*. Both bulk and single cell RNA-seq data indicated that following lineage switching, the activation of the new and silencing of the old lineage-selective gene program occurred in a transient manner while merging into one another. This suggests a potential trans-differentiation to occur without reactivation of a stable progenitor state.

### Some adipogenic committed cells maintain speciation in the absence of adipogenic signals

We have previously shown that enhancer activation during adipogenesis and osteogenesis of stromal cells differ, with adipogenesis relying on *de novo* enhancer activation while osteogenesis involves activation of pre-established enhancers (20). Based on network and single cell RNA-seq analysis, we further suggested a higher degree of self-maintaining regulatory circuits within adipogenic compared to osteogenic gene programs (20). Here, we detected mature lipid filled cells in cultures deprived from adipogenic inducers but not mineralized matrix in those deprived from osteogenic inducers. In line, scRNA-seq validated the presence of cells with a mature adipogenic gene signature that were refractory to switching from adipogenic to osteogenic stimulation (7d Ad + 4d Ob). In line with our observations, Schilling, T. et al. reported the abolishment of osteogenic gene expression (*ALPL* and *BGLAP*) upon switching to adipogenic inducers while adipogenic transcripts (*PPARG* and *LPL*) were still present upon switching to osteogenic inducers in human primary stromal cells (13). Likewise, Meyer, M. B. et al. reported that lineage switching in murine bone marrow-derived stromal cells from osteogenic to adipogenic inducers drastically decreased osteogenic transcripts (*Spp1* and *Runx2*) levels comparable to or lower than that of undifferentiated cells. In contrast, adipogenic transcript levels (*Pparg* and *Adipoq*) remained significantly higher after switching from adipogenic to osteogenic inducers compared to uninduced cells (26). Of note, our results might be biased with 7 days of initial differentiation being the more optimal time point for the establishment of the adipogenic transcriptional network compared to the osteogenic one. With the temporal stimulations we did, we suggest at least in some cells the transcriptional remodeling during adipogenesis to be a molecular switch or “point-of-no-return” in which transcriptional maturity is reached and maintained prior to phenotypic changes while osteogenic maturation occurs in a linear manner. Interestingly, the latter does not require continuous stimulation as alternating exposure to osteogenic and adipogenic inducers facilitated matrix mineralization.

### Induction of opposing lineage programs increased differentiation potential of stromal cells

Heterogeneity among primary stromal cells has been well known and varying osteogenic capacity has been reported between donor cells and for different subpopulations of individual donors (27, 28). Obtaining strong and reproducible lineage speciation of primary stromal cells is an unmet need for cell-based therapies and for the activation of endogenous progenitor cells to boost bone formation during regeneration and bone loss. Despite part of the interindividual differences in the osteogenic or adipogenic capacity of primary cells can be attributed to clinical and cellular features (24, 29), some primary cultures are unable to form mineralized matrix or lipid filled cells. Using hBMSC-TERT4 and primary cells we showed that preinduction towards the opposing lineage increases the adipogenic and osteogenic differentiation capacity *in vitro*, restores matrix mineralization in osteogenic non-responsive primary cells *in vitro*, and boosts bone formation of bone-forming competent cells in a xenograft assay. The first observation seemed to be also present in the studies by Schilling, T. et al. (13) and Meyer, M. B. et al. (26), in which mineralization was visually enhanced, however not quantified, when cells were first exposed to adipogenic inducers. Whether the lack of osteogenic potential in selected primary cultures is due to the absence of stimulating or presence of inhibiting signals is speculative, however, differentiation boosting effects of pre-stimulation with adipogenic inducers was persistent across different donors. Together with phenotype-switching experiments in which adipogenic or osteogenic primed human MSCs adopted alternative mesenchymal fates upon media change (14, 16), our data support the concept that prior lineage engagement can “train” stromal cells to respond more robustly to subsequent, even opposing, differentiation cues.

Beyond plasticity in cellular phenotype and gene expression, osteoblast–adipocyte interconversion also extends to the epigenetic level. Exposure to lineage-specific differentiation conditions drives the deposition of histone marks and lineage-defining transcription factor binding at osteoblast- and adipocyte-associated gene loci in a condition-dependent manner (26). We propose that osteogenic and adipogenic loci differ in their underlying epigenetic permissiveness to interconversion, reflecting a combination of progenitor-state priming and lineage-specific enhancer logic. Although differentiation cues can remodel enhancer landscapes, the absence of enhanced expression for a subset of genes upon interconversion suggests that only certain loci retain epigenetic responsiveness to opposing lineage signals, which might in part be due to more stable epigenetic mechanisms such as DNA methylation (30). Genes that remain epigenetically responsive to interconversion are likely to represent key regulatory or rate-limiting components of the differentiation program and may therefore disproportionately mediate the observed gains in osteogenic and adipogenic output. However, the increased differentiation potential upon pre-stimulation could also be associated with synchronization of the cells in terms of stochastic gene expression (31), induction of gene programs that are common between osteoblast and adipocyte differentiation, inhibition of stemness factors, promoting cell expansion, or due to sensitizing general cellular machineries responsible for transcription, translation, and signaling cascades which make cells more susceptible to respond to changes in extracellular stimuli.

### Transcriptional plasticity is not a singular event in vitro

We showed that the transcriptional plasticity of differentiating stromal cells *in vitro* was neither hampered by maturation of the cells nor repeated media switching which intuitively contrasts the hypothesis of terminally differentiated cells. We therefore propose that the niche and not the intrinsic cellular programs prevent lineage switching *in vivo*. Such a theory is supported by the observations of Clabaut et al. (9) showing osteoblasts to obtain an adipocyte like phenotype in co-culture settings *in vitro* and to express adipogenic markers in bone biopsy sections of elderly subjects. In line, our staining and single nuclei RNA-seq analysis strongly supported the notion of cells with adipogenic and osteogenic phenotypes and gene signatures being simultaneously present at least for a limited period of time. Similar observations at the level of osteogenic and adipogenic gene expression were made in primary human stromal cells sorted for high levels of ALP but also for ALP activity and lipid droplets as well as osteocalcin and PPARg protein levels when differentiating ALP-negative cells *in vitro* (22). This might explain why consecutive changes of differentiation media led to the presence of both lipid droplet accumulating adipocytes and matrix mineralizing osteoblasts. Despite we provided phenotypical staining and gene expression at single cell resolution, we did not track individual cells throughout consecutive rounds of media change. Being able to track single and multiple rounds of acquiring osteogenic and adipogenic phenotypes *in vitro* as well as *in vivo* would answer how directed trans-differentiation contributes to bone loss in conditions with increased marrow adipogenesis such as unloading (32), type I diabetes (33), osteoporosis (5), dietary intervention and radiation (34). Finally, since adipogenic stimulation did increase the osteogenic potential, it seems like a lack of bone-forming signals are of more importance for age-related bone loss than increased adipogenic signals.

## Materials and methods

### Cell culturing

hBMSC-TERT4 cells (23) or bio-banked primary stromal cells (24) were seeded in tissue culture flasks and expanded in culture medium consisting of minimum essential medium (MEM, 32561-094, Invitrogen) supplemented with 10% fetal bovine serum (FBS, 16000044, Life Technologies) and 1% penicillin/streptomycin (P/S, 15140-130, Invitrogen) and incubated at 37 °C with 5% CO_2_ and 95% humidity. hBMS-TERT4 cells were used at passages 43 to 50 and expanded at a split-ratio of 1:4 twice a week. Bio-banked primary cells were used at passage 2 to 4 and split 1:3 once a week. Culture medium was changed three times a week and the cells were either passaged or set out for experiments by trypsinization using 0.05% Trypsin-EDTA (25300062, Thermo Fisher Scientific) when reaching 80-90% confluency. Contamination with Mycoplasma was checked in the research unit every second month using either fresh conditioned media, or media stored at 4 °C until PCR test. All cultures reported here were negative of Mycoplasma.

### In vitro cell differentiation

#### Osteoblast differentiation

hBMSC-TERT4 cells and primary cells were seeded with a density of 30,000 and 20,000 cells/cm^2^ to achieve approximately 80% confluency the following day and expanded for 72 hours in culture medium. After 72 hours, culture medium was replaced with osteoblast induction medium consisting of MEM supplemented with 10% FBS, 1% P/S, 5mM ß-glycerophosphate (50020-100, Sigma-Aldrich), 10nM dexamethasone (D4902, Sigma-Aldrich), 50µg/mL 2-phosphate ascorbic acid (A4544, Sigma-Aldrich) and 10nM 1,25-vitamin D_3_ (D1530, Sigma-Aldrich). Media was refreshed every 2-3 days.

#### Adipocyte differentiation

hBMSC-TERT4 and primary cells were seeded with a density of 30 000 cells/cm^2^ to achieve approximately 100% confluency the following day and expanded for 72 hours in culture medium. After 72 hours, culture medium was replaced with adipocytic induction medium consisting of Dulbecco’s modified Eagle’s medium (DMEM, 41965062, Thermo Fisher Scientific) supplemented with 10% FBS, 1% P/S, 100nM dexamethasone, 3µg/mL insulin (I9278, Sigma-Aldrich), 1µM rosiglitazone (BRL, 71740, Cayman Chemicals) and 225µM 3-isobutyl-1-methylxanthine (IBMX, I5879-5G, Sigma-Aldrich). Media was refreshed every 2-3 days.

### Cellular assays for measuring osteogenic differentiation

#### Alizarin Red staining and quantification

Matrix mineralization was measured through Alizarin Red staining. Differentiated cells were washed with phosphate-buffered saline without Ca^2+^ and Mg^2+^ (PBS, 14190-169, Invitrogen), fixed with 77% ice-cold ethanol (51976, Sigma-Aldrich) at -20°C for 1 hour and washed with H_2_O. Fixed cells were stained with 40 mM Alizarin Red dye (A5533-25G, Sigma-Aldrich) dissolved in distilled H_2_O (pH=4.2) for 10 min with agitation at room temperature (RT) and washed with PBS. Stained cells were scanned, and images were captured with a magnification of 4x using an Olympus IX50 microscope. Alizarin Red staining was extracted for quantification by adding H_2_O with 20% methanol (65548, Sigma-Aldrich) and 10% acetic acid (20302236, VWR) to the cells and incubated for 10 min at RT with shaking. Absorbance was measured at 450 nm and 570 nm.

#### Alkaline Phosphatase (ALP) activity

Prior to measuring ALP activity, cell viability was measured through CellTiter-Blue assay according to manufacturer’s protocol (35) by replacing culture media with cell viability media (CellTiter-Blue (208657 (G8081), ILS Danmark) and culture medium, 1:5 respectively) and incubated at 37°C for 1 hour. Fluorescence intensity (579Ex/584Em) was measured with a FLUOstar Omega plate reader. For ALP activity, cells were washed twice with tris-buffered saline (TBS, (Lab44416.5000, Bie & Berntsen), pH=7,5), fixed in 3.7% formaldehyde (F1635, Sigma-Aldrich) – 90% ethanol for 30s and incubated with 1 mg/mL P-nitrophenylphosphate (dissolved in 50 mM NaHCO_3_ and 1 mM MgCl_2_ at pH=9.6, 71768-5G, Sigma-Aldrich) for 20 min. at 37°C. Absorbance was measured at 405 nm and ALP activity was normalized to cell viability.

#### ALP staining

Differentiated cells were washed with PBS, fixed in acetone (168014, ILS) / 10mM citrate buffer (168782, ILS) solution (3:2) at RT for 5 min and incubated in 0.2mg/mL Naphtol ((N6125-1G, Sigma-Aldrich) in H_2_O and N,N-Dimethylformamide (68-12-2, Sigma-Aldrich), 100:1) and 0.8mg/mL Fast red ((F8764, Sigma-Aldrich) in 0.1*M* Tris buffer) solution for 1 hour at RT. Cells were rinsed with H_2_O two times. Stained cells were scanned, and images were captured with a magnification of 4x using an Olympus IX50 microscope.

### Cellular assays for measuring adipocyte differentiation

#### Oil Red’O staining and quantification

Lipid accumulation was measured through Oil Red’O staining. Differentiated cells were washed with PBS, fixed with 4% paraformaldehyde (PFA, D383004, Hounisen) in PBS at RT for 10 min and washed with PBS. Fixed cells were stained with Oil Red’O solution (3:2 of 3mg/ml Oil Red’O (O0625, Sigma-Aldrich) in 100% isopropanol (I9516, Sigma-Aldrich) with H_2_O) for 1 hour at RT and rinsed with H_2_O. Images of lipid droplets were captured with a magnification of 10x using an Olympus IX50 microscope. Oil Red’O staining was extracted for quantification by adding isopropanol to the cells and incubated for 10 min at RT with agitation. Absorbance was measured at 500 nm.

### ELF^TM^ 97 Endogenous Phosphatase and Nile Red fluorescent labeling

Cells were stained with ELF 97 Endogenous Phosphatase Detection Kit (E6601, Thermo Fisher Scientific) and Nile Red (N3013, Sigma-Aldrich) according to manufacturer’s protocol. Briefly, cells were fixed in 4% PFA for 10 min at RT, washed twice with PBS, permeabilized in 0.2% Tween^®^ 20 (655205, Sigma-Aldrich) for 10 min and rinsed in PBS. Fixed and permeabilized cells were incubated in ELF phosphatase substrate diluted 20-fold in detection buffer for a few min at RT and development of fluorescent signal was monitored using a fluorescence microscope. Cells were washed in PBS and incubated with 5 µg/mL Nile Red dye (in PBS) for 10 min in the dark at RT. Fluorescence images of cells stained with Nile Red and ELF (485Ex/572Em and 345Ex/530Em, respectively) were taken with 4 and 10x magnification on a LEICA DFC300 FX microscope. To quantify fluorescent signal, we used ImageJ (1.53t) and converted images of individual fluorescent channels to 32-bit grey scales. Background was subtracted with a mask of 50.0 pixels. Adipocyte areas were manually defined using the red channel (NileRed) and the area was transferred on corresponding images of the blue channel (ELF). Pixel intensity was quantified for both channels at selected area.

### Ectopic bone formation

Assessment of bone-forming capacity using an *in vivo* transplantation assay has been previously described in detail (36). Shortly, we used 500,000 hBM-MSC-TERT4 cells first incubating with hydroxyapatite for 24 hours in syringes with growth media followed by maintaining the cells for 7 days in growth media or adipogenic differentiation media. We made 4 implants per mouse and compared bone formation to the same implantation sites in a paired manner. Implants were harvested after 8 weeks and fixed in 4% PFA for 24 hours followed by decalcification 0.5 M sodium format for 72 hours. Implants were paraffine embedded and sectioned at three different depths (100 µm apart). H & E stained sections were quantified for bone using ImageJ by marking areas of bone normalized to the total area of the implant. As a deviation from the published protocol, NOD.Cg-*Prkdc^scid^ Il2rg^tm1Sug^*/JicTac mice have been used instead of NOD/LtSz-*Prkdc*^scid^ mice. Ethical approval was given by the Animal Experimentation Council of the Ministry of Food, Agriculture and Fishing Denmark (License 2022-15-0201-01225).

### Bulk RNA-sequencing

RNA was extracted and purified from hBMSC-TERT4 cells using TRIzol (15596018, Invitrogen) and Econo Spin columns (Epoch Life Sciences, TX, USA) and unstranded RNA-seq libraries were constructed from 1 μg of total RNA according to manufactor’s instruction (TruSeq 2, Illumina). Libraries were sequenced on an Illumina Novaseq and reads were mapped to the human genome (GRCh38) using STAR (37), and tag counts were summarized at the gene level using HOMER (38) allowing only one read per position per length. Differential gene expression analysis and normalization were performed with DESeq2 (39).

### Enhancer dynamics based on DNase-seq and MED1 ChIP-seq

Processed DNase-seq and MED1 ChIP-seq data throughout osteoblast and adipocyte differentiation of hBM-MSCM-TERT4 cells (20) were merged with gene expression data. Each enhancer was linked to the closest RefSeqID based on annotation of enhancers using HOMER (38). Here we used normalized tag counts in undifferentiated cells and DEseq2 derived fold changes comparing 3 day of osteogenic or adipogenic differentiation to undifferentiated cells.

### Single cell and single nuclei RNA-sequencing

For single cell RNA-seq, hBMSC-TERT4 cells were trypsinized, spun down at 200 g and washed twice in culture media. Cells were separated using a 40 µm mesh, washed 3 times in PBS with 0.04% non-acetylated BSA, and cell numbers were adjusted to 0.7 x 10^6^ cells per ml followed by a final transfer through a 40 µm mesh. Using 10^4^ cells as input material single cell RNA-sequencing libraries were prepared according to the manufacturer’s instructions (10x Genomics). Cell gene count matrices were generated from raw sequencing data using zUMI pipeline (40) with human genome GRCh38 which yielded the following cell numbers: 2,597 undifferentiated; 4,434 for 7 days Ob; 8,630 for 7 days Ad; 7,251 for 7 days Ad + 4 day Ob; and 11,328 for 7 days Ob + 4 day Ad. Single cell data were merged and filtered for high quality droplets (mitochondrial percentage less than 10%, gene count higher than 200) using Seurat package (41) yielding a gene count matrix for 29,939 cells. Pseudotime and differential gene expression analysis was carried out using Monocle 2 (42).

For single nuclei RNA-seq, cells were trypsinized, centrifugated at 200 g for 5 min at 4°C, and supernatant was carefully removed without touching the upper layer. Cells were subjected to nuclei isolation following a protocol adjusted from Hauwaert et al. (43). Briefly, cells were dounced 5 times in nuclei isolation buffer consisting of 250 m*M* Sucrose (84097, Sigma-Aldrich), 10 m*M* HEPES (22320022, Thermo Fisher Scientific), 1.5 m*M* MgCl_2_ (M2670, Sigma-Aldrich), 10 m*M* KCl (P9541, Sigma-Aldrich), 0.001% IGEPAL CA-630 (I8896, Sigma-Aldrich), 0.2 m*M* DL-Dithiothreitol (43819, Sigma-Aldrich) and 0.5 U/mL RNase inhibitor (40,000 U/mL, M0314L, Bionordika) in DEPC-treated water. Nuclei were passed through a 30 µm cell strainer, centrifugated at 200 g for 10 min at 4°C and resuspended in nuclei resuspension buffer consisting of 1% BSA (A9418, Sigma-Aldrich) in PBS, 2 m*M* MgCl_2_ and 0.04 U/µL RNase inhibitor in PBS followed by a final transfer through a 30 µm cell strainer. Nuclei fixation followed by library preparation was carried out according to manufacturer’s instructions (Parse Biosciences Evercode WT, version 2). Raw sequencing reads processed with the Parse Biosciences Split Pipeline were mapped to the human genome (GRCh38) and yielded a gene count matrix of 5055 cells. Clustering was performed in Seurat (41). Doublet scores were calculated using scran (44). Quantification of previously defined osteoblast and adipocyte gene signatures (20) was done by summing scaled expression values.

For integrative PCA of bulk and single cell RNA-seq data we focused on genes with dynamic expression during osteogenic or adipogenic pseudotime (adjusted p < 0.001). Principal components were first computed from bulk RNA-seq data based on DESeq2 (39) log-transformed expression levels using the prcomp function in R with centering and scaling enabled. For the corresponding scRNA-seq samples, normalized expression values were averaged across pseudotime into 5 equally sized bins per condition and centered and scaled using the mean and standard deviation from the bulk RNA-seq matrix. These binned scRNA-seq profiles were then projected into the precomputed bulk PCA environment by multiplying the scaled expression matrix with the PCA rotation matrix (loadings), allowing direct comparison of bulk and single-cell trajectories in a common principal component space.

### Real time PCR (qRT-PCR)

RNA from hBMSC-TERT4 cells were extracted using TRIzol (15596018, Invitrogen) and isolated using Econo Spin columns (Epoch Life Sciences, TX, USA). Reverse-transcription of RNA into cDNA was achieved using the RevertAid H Minus First Strand cDNA Synthesis Kit (Thermo Scientific^TM^). Fast SYBR Green Master Mix (4385612, Applied Biosystems) and primers (specified below) were mixed to run quantitative Real-time PCR on an Applied Biosystems 7500 machine. Alpha-tubulin (*TUBA*) or TATA-binding protein (*TBP*) were used for normalization. Primer sequences for *ADIPOQ* (For: TGGGGGTGTCCTGGTACATGTGCAGAAAT; Rev: ACGCCTTTCATGACGCATTCCACCACC), *ALPL* (For: ACGTGGCTAAGAATGTCATC; Rev: CTGGTAGGCGATGTCCTTA), *TBP* (For: GCCCGAAACGCCGAATAT; Rev: CCTCATGATTACCGCAGCAAA) and *TUBA* (For: GAGGCTGACGCAGAATGCA; Rev: TCTGTGGCAATCCGGTTCA) were used.

### Statistical analyses

All statistical analysis were carried out in R, and unpaired student’s t-test was used for qPCR data and quantification of ALP activity, matrix mineralization, and lipid droplet formation in hBMSC-TERT4 cells. Paired student’s t-test was used for differentiation assays of primary cells. Statistics for sequencing-based methods were derived from Seurat, DEseq2, and Monocle 2. Replicates are indicated in the individual figures.

## Acknowledgements

Sequencing was carried out at the Villum Center for Bioanalytical Sciences, Functional Genomics & Metabolism Research Unit, University of Southern Denmark. We thank Ronni Nielsen for sequencing assistance. This work was supported by a grant from the Lundbeck Foundation (R335-2019-2195) and Novo Nordisk Foundation (NNF22OC0078257) to A.R. and a PhD scholarship from the Region of Southern Denmark to A.J.M.J.

## Author Contribution Statement (CRediT)

**Ali Jasim Mohammad Jamil:** Formal analysis, Investigation, Data curation, Visualization, Writing – original draft. **Mikkel Ørnfeldt Nørgård & Emilie Grupe**: Data curation. **Alexander Rauch:** Conceptualization, Funding acquisition, Supervision, Writing -review & editing

## Ethics Statement

Biobanked primary cells were previously obtained from orthopedic surgeries at Odense University Hospital with all subjects providing written consent and received oral and written information. The collection of human cells was conducted in accordance with the Declaration of Helsinki and approved by the Scientific Ethics Committee of the Region of Southern Denmark (project ID: S-20160084). Ethical approval for animal experiments was given by the Animal Experimentation Council of the Ministry of Food, Agriculture and Fishing Denmark (License 2022-15-0201-01225).

## Data Availability Statement

Raw sequencing data of hBMSC-TERT4 cells reported in this study have been deposited under the NCBI Gene Expression Omnibus: GSE288316. Scripts for data processing and visualization to recapitulate the analyses of sequencing-based data are available at the open science framework: https://osf.io/c84vs and at GitHub: https://github.com/drarauch/InVitroPlasticity.

